# Neural coding of isopentyl acetate alarm pheromone signals in the honey bee brain

**DOI:** 10.64898/2026.08.26.747191

**Authors:** Tao Lin, Brian H. Smith, Hong Lei

## Abstract

Alarm pheromone is a high-priority social signal in honey bees, yet direct evidence for how its major component, isopentyl acetate (IPA), is encoded in antennal lobe remains limited. Here, we combine intracellular recording, neuronal staining, and three-dimensional reconstruction to examine neural responses to IPA in the honey bee brain. Integrated analysis of antennal lobe neurons revealed clear but heterogeneous time-locked responses to IPA, which could be grouped into four temporal response motifs: fast transient, monophasic, biphasic excitation-inhibition, and delayed excitation-inhibition. A morphologically identified antennal lobe neuron exhibited a stable excitatory response characterized by short latency and prolonged elevated firing after stimulus onset. In a representative delayed-type antennal lobe neuron, response magnitude showed strong concentration dependence: peak amplitude and post-peak inhibition increased significantly with increasing IPA concentration, whereas peak latency remained largely unchanged. Repeated stimulation at an intermediate concentration produced comparatively modest effects, expressed mainly as attenuation of peak amplitude and a gradual delay in response timing. In addition to antennal lobe neurons, we identified two IPA-responsive protocerebral neurons. Together, these results provide direct single-neuron evidence that IPA is heterogeneously encoded in the honey bee antennal lobe.

## Introduction

Chemical communication is central to social organization in honey bee colonies. Among the many pheromonal signals used by bees, alarm pheromone is especially important because it links the detection of danger by individual workers to rapid colony-level defensive responses. The identification of isopentyl acetate, also known as isoamyl acetate, as an active component of the honey bee sting alarm pheromone provided the chemical foundation for studies of defensive communication in this species (Boch et al., 1962). Subsequent behavioral work showed that IPA is not merely an odor associated with stinging. Rather, it can recruit defensive workers, increase flight activity, and enhance responses to moving targets, making it one of the best-characterized chemical signals involved in honey bee defense (Wager and Breed, 2000).

The importance of IPA extends beyond its immediate role in eliciting defensive behavior. Several lines of evidence indicate that IPA can also alter behavioral state and response thresholds. European and Africanized honey bees differ in the intensity and duration of their responses to IPA or alarm-pheromone blends, suggesting that defensive responsiveness depends not only on the chemical stimulus itself but also on genetic background, baseline activity, and behavioral threshold (Collins et al., 1987). During worker development, responsiveness to IPA and 2-heptanone increases with age, and treatment with the juvenile hormone analog methoprene can accelerate the appearance of alarm responsiveness in young workers without changing their antennal electroantennogram responses (Robinson, 1987). Thus, the behavioral output evoked by alarm pheromone is unlikely to be explained by peripheral olfactory sensitivity alone. Response thresholds of interneurons and internal physiological state are also likely to be involved. Beyond triggering immediate defensive behavior, alarm pheromone can induce longer-lasting physiological and molecular changes in honey bees. Exposure to alarm pheromone has been shown to activate immediate-early gene expression and alter subsequent behavioral responses (Alaux and Robinson, 2007). IPA also affects learning and aversive responsiveness. Exposure to IPA or natural sting alarm pheromone impairs appetitive odor–sucrose learning, and this effect can last for many hours (Urlacher et al., 2010). Conversely, IPA can increase responsiveness to noxious electric shock, whereas 2-heptanone does not produce the same effect under comparable conditions (Rossi et al., 2018). Together, these findings suggest that IPA acts not only as a releaser of defensive behavior, but also as a state-modulating signal that can bias bees away from foraging, reward learning, and feeding-related responses and toward vigilance, defense, and risk avoidance.

Compared with this behavioral and physiological literature, the neural coding of IPA in the honey bee brain remains less completely understood. Calcium-imaging studies have shown that sting alarm pheromone components can evoke distinct activity patterns in the worker antennal lobe, the primary olfactory center of the insect brain. This view is consistent with broader observations that worker social pheromones, including IPA and 2-heptanone, are unlikely to be represented by a simple labeled-line system and may instead rely on across-glomerular and across-neuron activity patterns (Sandoz et al., 2007). Later work further showed that alarm-related compounds are represented in parallel antennal lobe output pathways and in the lateral horn, indicating that alarm-pheromone information is not restricted to a single primary olfactory route but is transmitted through distributed and higher-order olfactory circuits (Carcaud et al., 2015; Roussel et al., 2014; Carcaud et al., 2023).

Population imaging, however, cannot fully resolve the response diversity of individual neurons to IPA. This is an important gap. Alarm-pheromone coding may depend not only on which glomeruli or brain regions are activated, but also on spike timing, response duration, excitation–inhibition sequences, concentration-dependent gain, and changes across repeated stimulation. Intracellular studies in the honey bee have shown that antennal-lobe neurons can respond to odors with excitation, inhibition, or more complex temporal patterns, and that the same odor does not necessarily evoke the same response in different neurons (Müller et al., 2002; Galizia and Kimmerle, 2004). In addition, IPA can carry early odor-identity information through the relative latency order of glomerular responses, suggesting that timing itself is part of the neural representation of this alarm-related odor (Paoli et al., 2018). Understanding how IPA is processed as a socially urgent signal therefore requires direct analysis at the single-neuron level.

Here, we combined intracellular recording, neuronal staining, and three-dimensional reconstruction to examine how neurons in the honey bee brain respond to IPA. Rather than asking only whether IPA activates the antennal lobe, we focused on the temporal structure and physiological diversity of single-neuron responses, their dependence on stimulus strength and repeated stimulation, and the possibility that IPA-responsive activity is represented beyond the primary olfactory center. This approach provides direct single-neuron evidence for how a socially important alarm signal is encoded in the honey bee brain and helps bridge previous behavioral, population-imaging, and theoretical studies of pheromone processing.

## Materials and methods

### Experimental animals and preparation

Foraging worker honey bees, *Apis mellifera*, were used in this study. Bees were collected from multiple colonies maintained at the Arizona State University apiary. To avoid sampling guard bees, only outbound foragers leaving the hive entrance were captured with a glass vial, whereas bees standing or patrolling at the entrance were not used. Captured bees were transferred to perforated sample bottles and brought to the laboratory.

Bees were anesthetized on ice for 3–5 min. The head was fixed on a small platform with low-melting wax, and molten eicosane (melting point 36.8 °C) was applied to the base of the scape and to the pedicel–flagellum joint to stabilize the antennae. A small window was opened in the head capsule between the compound eyes, and fat body, glands, and tracheae overlying the brain were removed carefully while avoiding damage to the antennal nerves. The perineural sheath covering the brain was then removed to facilitate electrode penetration. To minimize brain movement caused by proboscis muscles, the clypeus was opened and the muscles and tissues connecting the proboscis and tentorium were removed. The head capsule was continuously superfused with saline containing (in mM): 130 NaCl, 6 KCl, 4 MgCl_2_, 5 CaCl_2_, 160 sucrose, 25 glucose, and 10 HEPES, pH 6.7 (Galizia and Vetter, 2005).

### Intracellular recording and staining

Glass capillaries (Sutter Instruments, BF100-78-10) were pulled with a laser puller (Sutter Instruments, P-2000) to produce intracellular microelectrodes. When filled, electrode resistance ranged from 100 to 300 MΩ. Electrode tips were backfilled by capillary action with Lucifer Yellow solution, and the shaft was subsequently filled with lithium chloride using a Microfil needle (34 Gauge; World Precision Instruments, Sarasota, FL, USA), taking care to avoid air bubbles. The prepared electrode was mounted in a MEH2R electrode holder (World Precision Instruments) connected to an AxoClamp 2B amplifier.

The headstage was mounted on a Leica micromanipulator positioned on a TMC vibration-isolation table. A reference electrode was inserted into the compound eye. Under a stereomicroscope equipped with an Olympus DF PLAPO 1× objective and Olympus WHSZ10×-H/22 eyepieces, the recording electrode was advanced into the target brain region. Impalement was achieved by brief pulsed microcurrent buzzing followed by fine adjustment to stabilize the recording. Signals were digitized at 25 kHz with an NI USB-6363 A/D converter (National Instruments, Austin, TX, USA) and recorded using the custom acquisition program SpikeHuond (v1.2).

Recording sites in the antennal lobe were guided by regions previously shown by calcium imaging to respond strongly to IPA and related alarm-pheromone stimuli (Wang et al., 2008; Sandoz et al., 2007). Protocerebral recordings were targeted mainly to the region surrounding the mushroom body α-lobe, based on the morphology of feedback neurons described by Grünewald (1999). Only recordings with stable spike waveforms and clear stimulus-locked responses were included in subsequent analyses.

### Odor stimulation

Odor stimuli were delivered using paper strips (Whatman No. 5) loaded with 2 μl odor solution and inserted into 0.5-ml glass syringes. Airflow through the syringe was controlled by a three-way solenoid valve, and both NC and NO outlets were connected to an L-shaped glass tube carrying a constant airstream of approximately 1 L min^-1. Stimulus trains consisted of 5 or 10 pulses, each 500 ms in duration, with an inter-pulse interval of 4.5 s. The delivery line was flushed for at least 35 s before switching to a different stimulus concentration.

For concentration-dependent analyses, IPA was tested at four concentrations (0.001, 0.01, 0.1, and 1, v/v in mineral oil) prepared as mineral oil dilutions and presented in randomized order across preparations. For repeated-stimulation analysis at 0.1, responses were analyzed across 18 trials; early trials were defined as the first 9 trials and late trials as the last 9 trials.

### Data analysis and statistics

Spike timestamps were extracted with a third-party Matlab program SpikeHuond, and all subsequent analyses were performed in MATLAB using custom-written scripts. Spikes were binned at 50 ms and converted to firing rate (Hz). For each trial, responses were baseline-corrected by subtracting the mean firing rate during the 0.5 s prestimulus period. Response parameters were quantified within the poststimulus window used for each analysis (0–2.5 s for the analyses of eight recorded antennal lobe neurons and representative-neuron analyses in Figs. 1, 2, and 4, and 0–3.0 s for the concentration- and repeated-stimulation analyses in Fig. 3).

**Figure 1.**
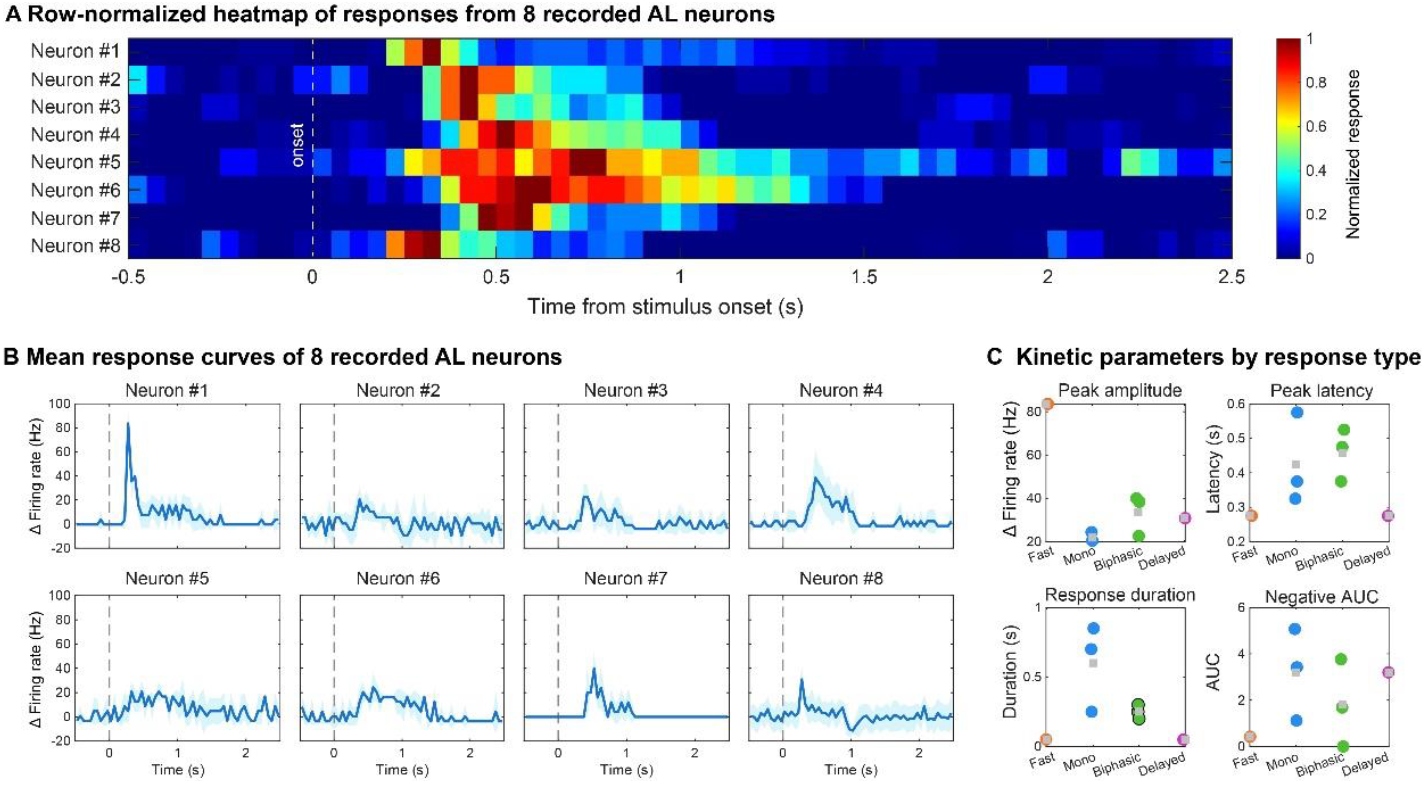
Response patterns and response-motif classification of antennal lobe neurons in response to IPA stimulation. (A) Row-normalized heat map showing baseline-corrected responses of eight recorded antennal lobe neurons aligned to stimulus onset. Each row represents one neuron, and color intensity is normalized to the maximum post-stimulus response of that neuron. Warmer colors indicate stronger responses. The dashed line marks stimulus onset (0 s) (i.e. Solenoid valve opening). (B) Averaged response curves for the same set of neurons. The blue trace shows the mean change in firing rate relative to baseline, and the shaded region indicates trial-to-trial standard deviation (±SD). The dashed line marks stimulus onset. (C) Descriptive summary of four kinetic parameters for the eight neurons, including peak amplitude, peak latency, response duration, and negative AUC (area under curve), shown to illustrate the basis for grouping them into four response motifs. Colored circles represent individual neurons, and gray squares indicate the mean within each response motif.

**Figure 2.**
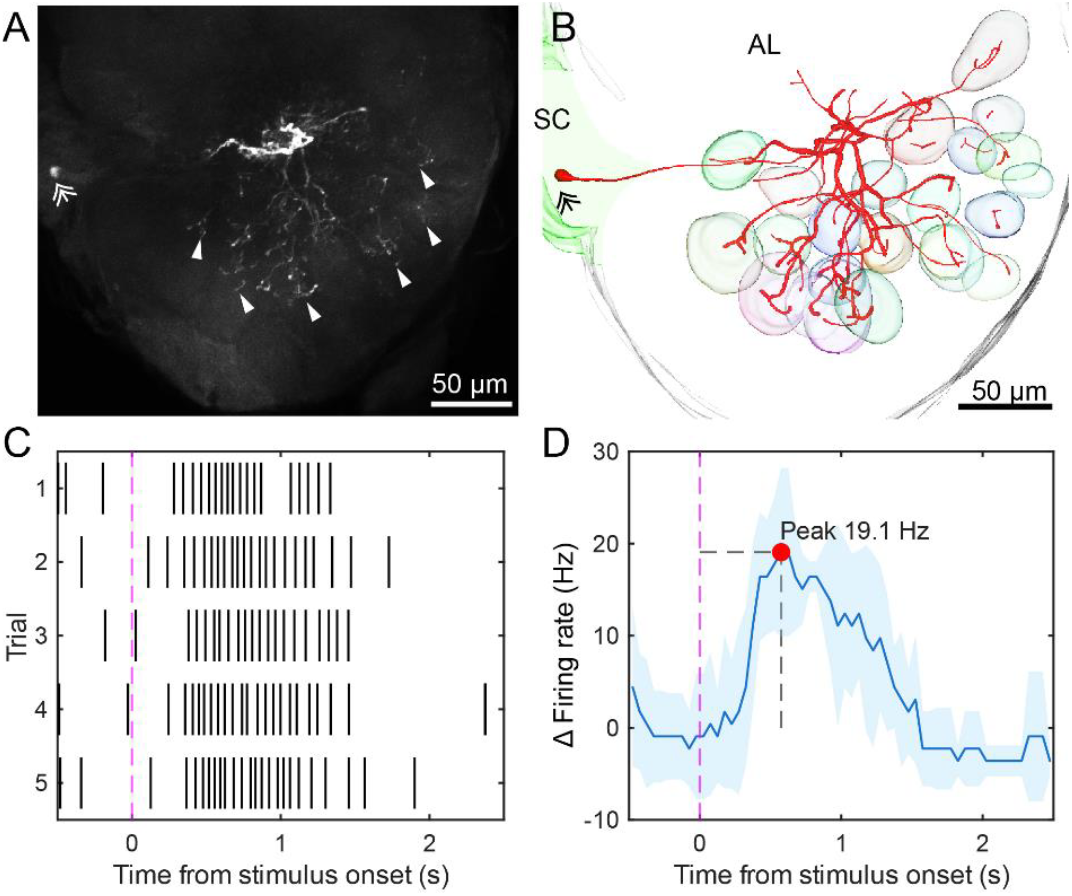
Morphological reconstruction and electrophysiological response properties of a representative IPA-responsive antennal lobe neuron. (A) Confocal image of the successfully labeled neuron. Double arrowheads indicate the soma, and single arrowheads indicate terminal arborizations within antennal lobe neuropil. (B) Three-dimensional reconstruction of the same neuron, showing its branching pattern within the antennal lobe (AL); SC indicates the soma cluster. (C) Spike raster plot of the neuron during repeated IPA stimulation. The magenta dashed line marks stimulus onset. (D) Mean change in firing rate aligned to stimulus onset. The blue trace indicates the mean response, the light blue shading indicates trial-to-trial variability, the red dot marks the response peak, and the gray dashed lines indicate peak amplitude and peak latency. Kinetic values shown in panel D were extracted from the smoothed trial-averaged response curve for visualization. Scale bars in A and B = 50 μm.

**Figure 3.**
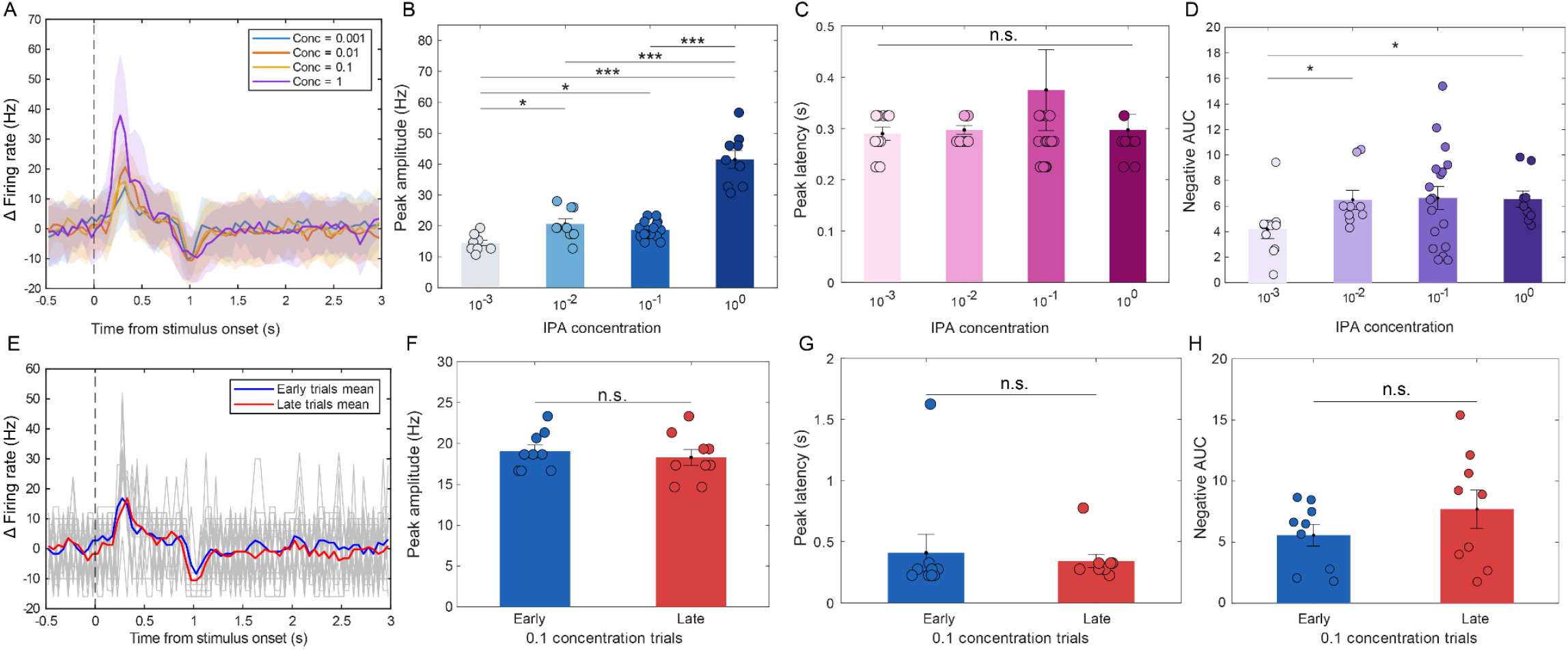
Concentration-dependent and repeated-stimulation response properties of a representative delayed-type antennal lobe neuron (Neuron #8). (A) Mean hange in firing rate at different IPA concentrations. Traces represent baseline-corrected average responses, the shaded regions indicate trial-to-trial variability, and the dashed ne marks stimulus onset. (B) Comparison of peak amplitude across IPA concentrations. (C) Comparison of peak latency across IPA concentrations. (D) Comparison of ost-peak inhibition strength (negative AUC) across IPA concentrations. (E) Mean firing-rate responses during repeated stimulation at concentration 0.1. Gray traces show l individual trials, the blue trace shows the mean of early trials (first 9 trials), the red trace shows the mean of late trials (last 9 trials), and the dashed line marks stimulus nset. (F) Comparison of peak amplitude between early and late trials at concentration 0.1. (G) Comparison of peak latency between early and late trials at concentration 0.1. H) Comparison of post-peak inhibition strength between early and late trials at concentration 0.1. Bars indicate means, error bars indicate SEM, and circles indicate individual ial values. Horizontal lines and asterisks denote significant group differences; “n.s.” indicates a non-significant difference.

**Figure 4.**
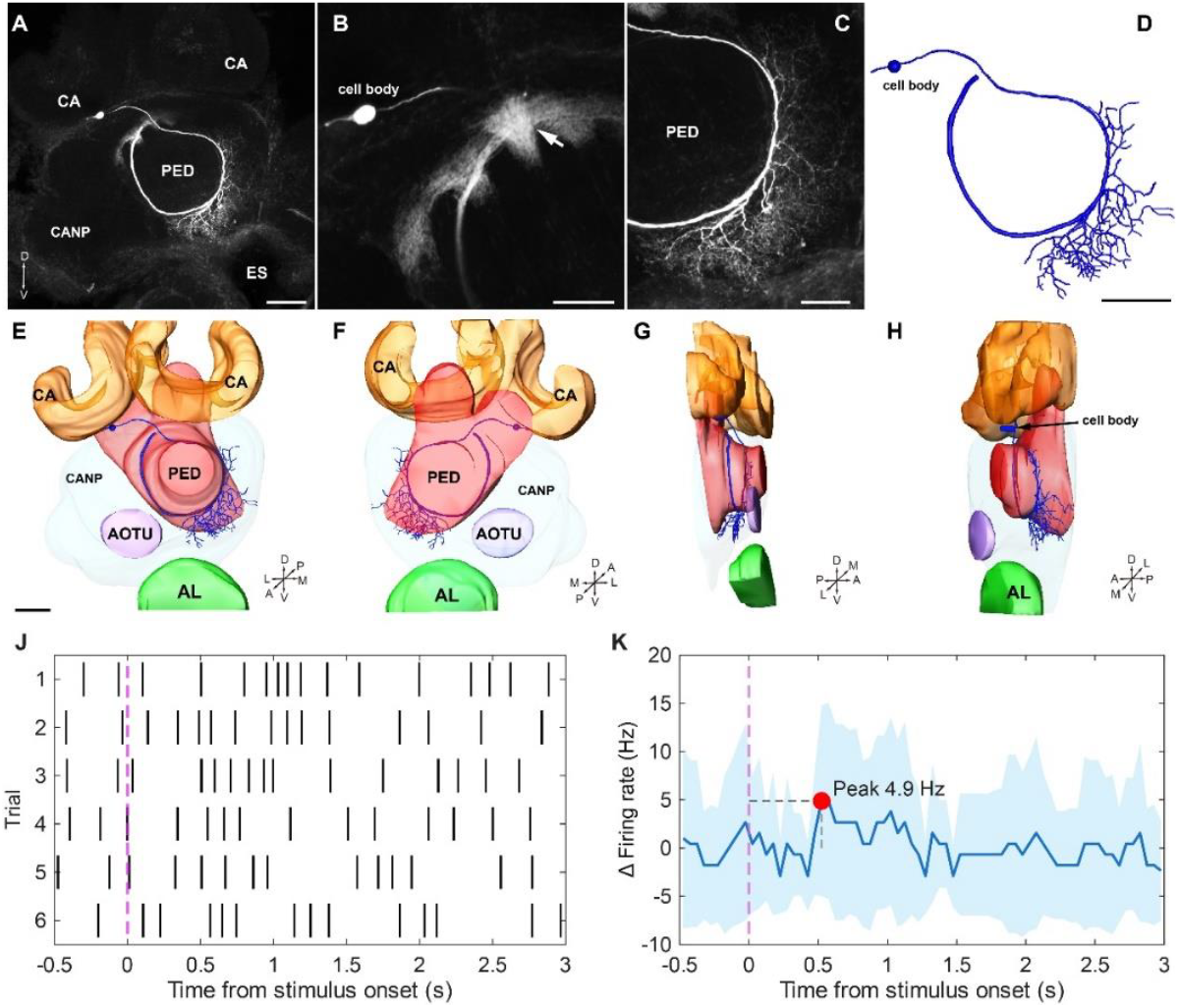
Morphological reconstruction and electrophysiological response properties of an IPA-responsive protocerebral neuron. (A–C) Confocal images of the successfully labeled neuron, showing that the soma is located near the calyx (CA) and that the neurites are distributed around the pedunculus (PED) and adjacent protocerebral regions; in B, the arrow indicates a major branching region in the PED. (D) Skeleton reconstruction of the same neuron. (E–H) Three-dimensional reconstructions from different viewing angles showing the spatial relationships of the neuron with the calyx (CA), pedunculus (PED), anterior optic tubercle (AOTU), antennal lobe (AL), and surrounding protocerebral neuropils. (J) Spike raster plot during repeated IPA stimulation. The magenta dashed line marks stimulus onset. (K) Mean change in firing rate aligned to stimulus onset. The blue trace indicates the mean response, the light blue shading indicates trial-to-trial variability, the red dot marks the response peak, and the gray dashed lines indicate peak amplitude and peak latency.

For the descriptive analyses of the eight recorded antennal lobe neurons, trial-averaged baseline-corrected responses were used to generate heat maps and mean response curves. In the heat map, each neuron’s response was row-normalized to its own maximum poststimulus response. For representative single-neuron display panels, raster plots and mean firing-rate traces were generated from the same baseline-corrected data. In representative averaged-response panels, a 3-bin moving average was applied for visualization of response kinetics, and the kinetic parameters shown in those panels were extracted from the smoothed trial-averaged response curve. Representative kinetic values summarized for individual antennal lobe neurons were calculated from unsmoothed trial-averaged baseline-corrected responses.

Peak amplitude was defined as the maximum poststimulus increase in firing rate. Peak latency was defined as the time from stimulus onset to the response peak. Response duration was calculated as the cumulative time during which the response remained at or above 50% of the peak amplitude. Post-peak inhibition was quantified as the negative area under the curve (negative AUC), calculated from the portion of the baseline-corrected response falling below zero during the poststimulus period. Based on the temporal profiles of averaged responses, antennal lobe neurons were classified as fast transient, monophasic, biphasic excitation-inhibition, or delayed excitation-inhibition. The comparisons among response types shown in Fig. 1C were descriptive and were used to summarize kinetic differences across neurons, rather than to perform formal inferential statistical tests.

For the representative delayed-type antennal lobe neuron, associations between response parameters and log10-transformed IPA concentration were tested using Spearman rank correlation. Differences among concentration groups were evaluated with Kruskal-Wallis tests followed, when appropriate, by Bonferroni-corrected pairwise rank-sum tests. For repeated-stimulation analysis at 0.1, changes across trials were assessed using Spearman rank correlation with trial number, and early versus late trials were compared using rank-sum tests. Bar plots show mean ± SEM, whereas shaded regions in averaged response curves indicate trial-to-trial variability (±SD). Statistical significance was set at *P* < 0.05.

### Image acquisition and 3D reconstruction of neurons

After physiological characterization, neurons were filled by injecting hyperpolarizing current to allow Lucifer Yellow to diffuse into the recorded neuron, after which the electrode was carefully withdrawn. The brain was then dissected from the head capsule, rinsed in saline, and kept in a humidified dark chamber for 1–2 h to allow the tracer to diffuse throughout the neuron. Brains were fixed overnight (9–10 h) in 4% paraformaldehyde at 4 °C, rinsed in PBS four times for 15 min each, dehydrated through an ethanol series (30%, 50%, 70%, 95%, and 100%), and then cleared in methyl salicylate. Cleared brains were stored in methyl salicylate until imaging. For confocal observation, each brain was placed in a well of a custom-made aluminum slide, immersed in methyl salicylate, and imaged directly without permanent mounting.

Samples were imaged with a Zeiss LSM710/780 confocal microscope (Jena, Germany). The neuronal tracer was excited with a 488-nm laser line, and image stacks were acquired at a z-step of 1–3 μm and a resolution of 1024 × 1024 pixels. Major brain neuropils were segmented manually in Amira

5.3.2 using the LabelField module, and labeled neurons were reconstructed in three dimensions using the Skeleton Tree tool. Neuropil orientation followed the insect body-axis convention described by Ito et al. (2014).

### Sample size and reporting

A total of 36 bees were used in this study. Across these preparations, 45 intracellular recording attempts were made, of which 10 yielded stable recordings that met the inclusion criteria for analysis. Among these, 3 neurons were successfully stained and reconstructed. Figure 1 and Table S1 summarize responses from eight recorded antennal lobe neurons. One morphologically identified antennal lobe neuron (Neuron #6) was used for the representative structural and electrophysiological analysis shown in Fig. 2. One delayed-type antennal lobe neuron (Neuron #8) was used for the concentration-dependent and repeated-stimulation analyses shown in Fig. 3. Figures 4 and 5 each present one successfully labeled protocerebral neuron that showed a clear response to IPA.

**Figure 5.**
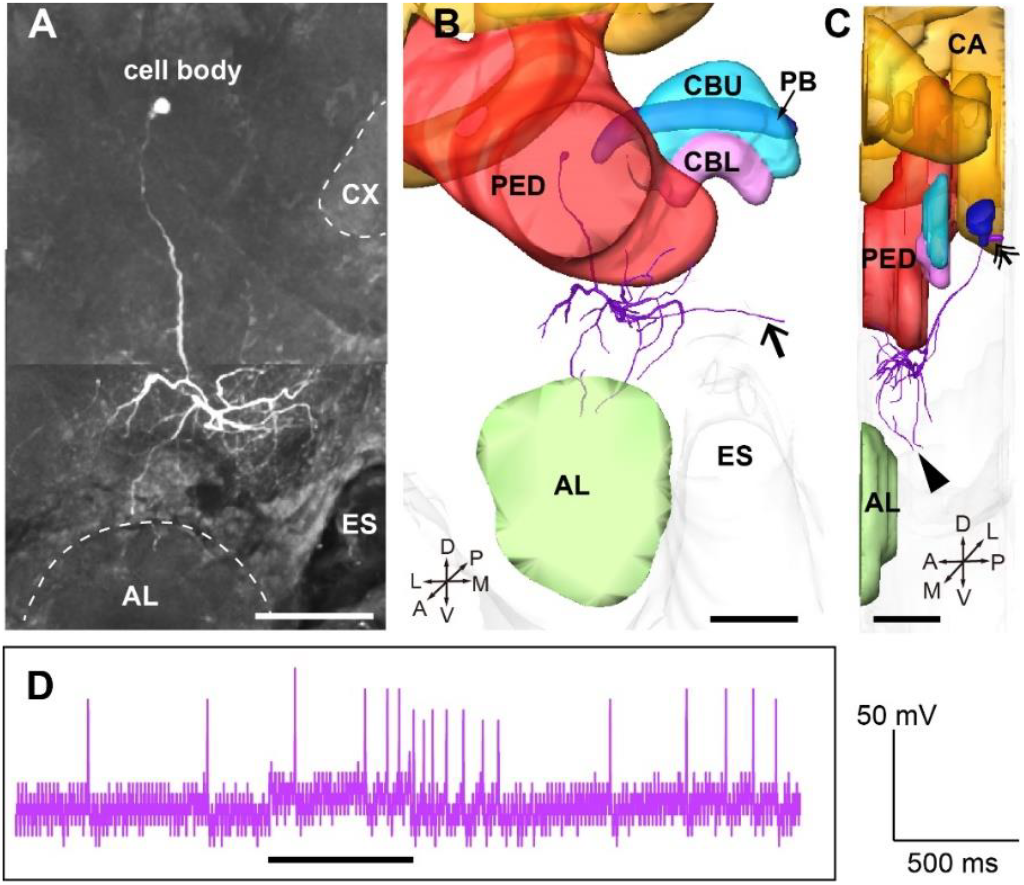
Morphological features and excitatory response of a protocerebral neuron to IPA. (A) Confocal section projection showing the location of the neuron in the protocerebrum. The soma is located ventral to the calyx (CA), and its dendrites are distributed dorsal to the antennal lobe. (B–C) Three-dimensional reconstruction of the neuron using Amira software to show its location in the brain. B is a frontal view and C is a medial view; double arrowheads indicate the soma. The axon arises from a cell body located in the posterior part of the brain and branches mainly between the antennal lobe and the mushroom body lobes. Some branches project through the esophageal commissure to the contralateral side (arrows), whereas others enter the medial lobe but not the antennal lobe (arrowheads). (D) Neuronal activity recorded during IPA stimulation of the antenna, demonstrating a clear excitatory response to IPA.

## Results

### 3.1 Response diversity and kinetic features of IPA-evoked responses

Integrated analysis of eight recorded antennal lobe (AL) neurons showed that isopentyl acetate (IPA) elicited clear, time-locked responses, although the response profiles varied markedly among neurons (Fig. 1). The row-normalized heat map showed that the principal response of most neurons occurred approximately 0.3–1.0 s after stimulus onset (defined here as solenoid valve opening), but substantial differences were evident in response strength, peak timing, and response duration (Fig. 1A). Averaged response curves further demonstrated that AL neurons did not respond to IPA in a uniform manner (Fig. 1B). Neuron #1 exhibited a high-amplitude, short-latency fast transient increase. Neurons #2, #5, and #6 showed sustained monophasic excitation. Neurons #3, #4, and #7 displayed biphasic excitation-inhibition profiles, characterized by a rapid decline after the excitatory peak together with a distinct inhibitory component. Neuron #8 showed a delayed response with relatively low amplitude.

Overall, the eight neurons differed in peak amplitude, peak latency, response duration, and the strength of post-peak suppression. Based on their temporal response profiles, they could be grouped into four response motifs: fast transient, monophasic, biphasic excitation-inhibition, and delayed (Fig. 1C, Table S1). Figure 1C provides a descriptive summary of four kinetic parameters across the eight neurons and illustrates the basis for grouping them into these four response motifs. In particular, the fast transient neuron showed the largest peak amplitude and the shortest response duration, monophasic neurons tended to show more prolonged responses, biphasic excitation-inhibition neurons exhibited stronger post-peak suppression, and the delayed neuron was characterized by a later peak time and intermediate response amplitude. Together, these observations indicate that IPA responses in the recorded AL neurons were heterogeneous and could be described by multiple temporal response motifs with distinct kinetic features.

### 3.2 Morphological reconstruction and electrophysiological response properties of a representative IPA-responsive antennal lobe local interneuron

Among all recorded neurons, one antennal lobe neuron showing a clear response to IPA was successfully labeled morphologically (Neuron #6; Fig. 2). Confocal imaging showed that its soma was located in the ventrolateral soma cluster of the antennal lobe, with neurites extending into the AL (Fig. 2A). Three-dimensional reconstruction further revealed that its terminal branches were distributed mainly within ventromedial glomerular regions of the AL, forming an extensive local arborization pattern (Fig. 2B).

Electrophysiological recordings showed that this neuron exhibited a reliable stimulus-locked excitatory response to IPA across repeated trials (Fig. 2C). Across five repeated stimulus presentations, a clear increase in firing was observed after stimulus onset in every trial, indicating that the neuron was consistently activated by IPA. Although response magnitude varied somewhat across trials, the temporal response profile was highly consistent, with firing rising rapidly after stimulus onset, remaining elevated for several hundred milliseconds, and then decaying gradually toward baseline. Analysis of the averaged response curve used for display showed a response peak at approximately 0.575 s after stimulus onset, with a peak amplitude of 19.07 Hz, a response duration of 0.90 s, and a negative AUC of 2.580 (Fig. 2D). These values were extracted from the smoothed averaged trace for visualization and therefore differ slightly from the representative kinetic values summarized in Table S1. Overall, this neuron exhibited a relatively large-amplitude, short-latency, and prolonged excitatory response to IPA, accompanied by a modest inhibitory component after the excitatory peak.

### 3.3 Concentration-dependent and repeated-stimulation response properties of a representative delayed-type antennal lobe neuron

In the representative delayed-type antennal lobe neuron, Neuron #8, IPA-evoked responses showed a pronounced concentration dependence (Fig. 3). Averaged response curves indicated that, as IPA concentration increased, the excitatory peak became progressively larger and the post-peak inhibitory component became more prominent (Fig. 3A). Peak amplitude was positively correlated with log10-transformed IPA concentration (Spearman, *ϱ* = 0.783, *P* = 1.32 × 10 ^ -10 , and the overall effect of concentration was significant (Kruskal–Wallis, *H* = 29.37, *P* = 1.87 × 10 ^ -6; Fig. 3B). Post hoc comparisons further showed that peak amplitude did not differ between 0.001 and 0.01 (*P* = 1.000, Bonferroni-corrected), but was significantly greater at 0.1 and 1 than at 0.001 (*P* = 0.0167 and 0.00013, respectively) and at 0.01 (*P* = 0.0496 and 0.000247, respectively). In addition, the response at concentration 1 was significantly larger than that at 0.1 (*P* = 0.000333), indicating a progressive enhancement of excitatory output with increasing stimulus intensity. In contrast, peak latency did not vary significantly across concentrations (Spearman, ρ = 0.241, *P* = 0.107; Kruskal–Wallis, *H* = 4.11, *P* = 0.250; Fig. 3C), suggesting that stimulus concentration primarily modulated response magnitude rather than response timing. Negative AUC also showed a significant concentration dependence (Spearman, ρ = 0.363, *P* = 0.0131; Kruskal–Wallis, *H* = 12.22, *P* = 0.00668; Fig. 3D). Specifically, the inhibitory component at the lowest concentration (0.001) was significantly weaker than that at 0.01, 0.1, and 1 (*P* = 0.00416, 0.0463, and 0.0153, respectively), whereas no significant differences were detected among 0.01, 0.1, and 1 (all *P* = 1.000). These results indicate that post-peak inhibition is weak at very low stimulus strength, but increases rapidly and then reaches a plateau across the higher concentration range. To our knowledge, previous honeybee antennal lobe studies have reported inhibitory and excitation-inhibition responses, but have not described this specific concentration-dependent profile of post-peak inhibition in a single IPA-responsive neuron.

Repeated stimulation at concentration 0.1 also modulated the response dynamics of this neuron, although the effect was weaker than that of concentration (Fig. 3E). The averaged response profiles of early and late trials were broadly similar, but late trials tended to exhibit a slightly smaller excitatory peak and a modest delay in the timing of the peak. Consistent with this trend, peak amplitude showed a negative relationship with trial number that did not reach significance (Spearman, ρ = -0.383, *P* = 0.117), whereas the comparison between early and late trials revealed a significant reduction in peak amplitude in late trials (rank-sum test, *P* = 0.0406; Fig. 3F). Peak latency increased significantly with trial number (Spearman, ρ = 0.514, *P* = 0.0291), indicating that responses became progressively slower over repeated stimulation, although the direct comparison between early and late trials was not significant (rank-sum test, *P* = 0.529; Fig. 3G). By contrast, negative AUC showed neither a significant association with trial number (Spearman, ρ = 0.148, *P* = 0.558) nor a significant difference between early and late trials (rank-sum test, *P* = 0.475; Fig. 3H). Thus, coding of IPA by Neuron #8 was dominated by a strong concentration-dependent component, whereas repeated-stimulation effects were comparatively modest and were expressed mainly as attenuation of excitatory peak amplitude and a gradual delay in response timing rather than a major reorganization of the overall response pattern.

### 3.4 Morphological reconstruction and electrophysiological response properties of an IPA-responsive protocerebral neuron

To determine whether IPA-related activity extends beyond the antennal lobe, we recorded and labeled one neuron located in the protocerebrum that responded to IPA (Fig. 4). Confocal imaging showed that its soma was located near the calyx (CA), with the main neurite extending into deeper protocerebral regions and forming a relatively rich arborization around and adjacent to the pedunculus (PED) (Fig. 4A–C). Skeleton reconstruction and three-dimensional visualization further showed that its projections were concentrated around the pedunculus and were spatially associated with the calyx, anterior optic tubercle (AOTU), antennal lobe (AL), and surrounding protocerebral neuropils (Fig. 4D– H).

Electrophysiological recordings showed a clear time-locked change in spiking during repeated IPA stimulation (Fig. 4J). Although there was some variability across trials, the overall firing rate increased after stimulus onset. The averaged response curve showed a response peak at approximately 0.525 s after stimulus onset, with a peak amplitude of about 4.89 Hz, a response duration of 0.500 s, and a negative AUC of 1.817 (Fig. 4K). Compared with strongly responsive antennal lobe neurons, this protocerebral neuron showed a smaller response amplitude and a shorter response duration, but it nevertheless displayed a stable and clearly detectable stimulus-related dynamic change together with a modest negative component.

### 3.5 Morphological features and excitatory response of a protocerebral neuron to IPA

We next examined a second protocerebral neuron with a more extensive projection pattern that also showed a clear response to IPA (Fig. 5). The soma of this neuron was located in the dorsal protocerebrum, and the primary neurite extended ventrally, forming relatively dense terminal arborizations in the ventral protocerebrum (Fig. 5A). Three-dimensional reconstruction further showed that its branches were concentrated near the pedunculus (PED) and were located in the vicinity of the upper division of the central body (CBU), protocerebral bridge (PB), lower division of the central body (CBL), calyx (CA), and antennal lobe (AL). (Fig. 5B, C). This neuron formed distinct local branches around the pedunculus while extending ventrally into lower protocerebral regions.

Electrophysiological recording showed a clear excitatory response to IPA (Fig. 5D). During the stimulus period, neuronal activity increased markedly, as reflected by an elevated spike rate and a larger number of spike events. In contrast, baseline activity before stimulation was relatively low and stable. After stimulus onset, the neuron rapidly entered a high-activity state and maintained elevated firing for a period of time, indicating that this protocerebral neuron could be effectively activated by IPA.

## Discussion

The main finding of this study is that IPA did not evoke a single, uniform excitatory response in honey bee antennal lobe neurons. Instead, IPA responses showed clear temporal diversity. Based on the averaged response profiles, the eight antennal lobe neurons could be grouped into four response types: fast transient, monophasic, biphasic excitation–inhibition, and delayed excitation–inhibition. This result is consistent with the broader view that social pheromones in worker honey bees are not necessarily processed through a strict labeled-line system, but may instead be represented by distributed activity across neurons that differ in response timing, duration, and inhibitory structure (Sandoz et al., 2007). In this respect, the coding of IPA may differ from the more specialized organization often described for insect sex pheromone systems. In many moths, sex pheromone information is detected by highly selective receptor channels and processed through anatomically specialized pathways that include the macroglomerular complex and male-specific pheromone-selective projection neurons (Christensen and Hildebrand, 1987; Hansson et al., 1991). More broadly, such sex pheromone pathways have often been considered examples of segregated or relatively dedicated parallel olfactory processing streams (Galizia and Rössler, 2010). At the same time, sex pheromone coding is not always purely labeled-line, and work in moths has shown that both labeled-line and across-fiber processing can contribute to central pheromone representation (Jarriault et al., 2009). Against this background, the present results suggest that the alarm pheromone component IPA in worker honey bees is represented in a more distributed and temporally diverse manner at the level of antennal lobe neurons, rather than through a single highly specialized pheromone channel.

This temporal diversity may be relevant to the behavioral role of alarm pheromone. Alarm signals are typically encountered during predator attack, stinging, nest disturbance, or danger at a food source, and they require rapid changes in individual and colony-level defensive state (Nouvian et al., 2016; Wang and Tan, 2019). In this context, fast transient neurons may contribute to the rapid detection of an alarm cue, whereas monophasic neurons with more sustained excitation may help maintain a short-lived alarm state after stimulus onset. Biphasic and delayed excitation–inhibition responses may contribute to signal termination, temporal contrast, or the prevention of excessive excitation. These functional interpretations remain tentative and will require behavioral or circuit-level tests. Nevertheless, they are compatible with the idea that timing is an important part of odor coding in the honey bee brain. Paoli et al. showed that odor identity can be represented by the relative latency order of glomerular responses, and IPA was among the odorants showing a stable latency structure (Paoli et al., 2018). The differences observed here in peak latency, response duration, and post-peak inhibition further support the view that temporal dynamics contribute to the central representation of IPA.

The morphology and physiology of the representative antennal lobe neuron, Neuron #6, further show that IPA can reliably activate specific antennal lobe neurons. This neuron had its soma in the ventrolateral soma cluster of the antennal lobe, with arborizations mainly distributed in a ventromedial region, and it responded to IPA with a relatively short latency and a stable, sustained excitatory response. This general location is broadly compatible with the medial antennal lobe activity reported for IPA in calcium-imaging studies (Wang et al., 2008). However, because the specific glomerular identity of this neuron was not mapped onto a standardized antennal lobe atlas, we cannot determine whether it corresponds to a previously identified IPA-responsive glomerulus. Galizia and Kimmerle showed that the odor response of an individual antennal lobe neuron is closely related to the glomerulus it innervates, but is also shaped by local network interactions and neuronal type (Galizia and Kimmerle, 2004). Thus, based on its morphology and physiology, Neuron #6 is clearly a local interneuron in the antennal lobe that showed some of the typical LN responses - fast, relatively small in firing-rate amplitude, reliable responses across repeated trials – as described in Meyer et al. (2013).

The delayed-type neuron, Neuron #8, provided a useful single-neuron example of concentration-dependent IPA coding. As IPA concentration increased, both the excitatory peak and the post-peak inhibitory component increased, whereas peak latency remained relatively stable. This suggests that, in this neuron, IPA concentration was represented mainly through response magnitude and inhibitory strength rather than through a marked shift in response timing. Behavioral studies have shown that IPA effects are strongly dose- and state-dependent. Depending on dose and experimental context, IPA can enhance defensive or aversive responsiveness, induce stress analgesia, suppress foraging and dancing, or impair appetitive learning (Núñez et al., 1997; Balderrama et al., 2002; Urlacher et al., 2010; Gong et al., 2017; Rossi et al., 2018). Against this background, the concentration dependence of Neuron #8 may represent one form of intensity coding in the antennal lobe. A stronger excitatory peak at higher concentrations could provide downstream circuits with a stronger alarm-related input, whereas the enhanced post-peak inhibition may reflect local network regulation following stronger stimulation. Intracellular studies have shown that honey bee antennal lobe neurons can respond to odors with excitation, inhibition, or combined excitation–inhibition patterns (Müller et al., 2002; Galizia and Kimmerle, 2004). The parallel increase in excitation and inhibition observed here fits within this general framework, although the specific circuit mechanisms remain to be determined.

A related possibility is that the temporal structure of this neuron’s response may also carry information about stimulus timing. In Neuron #8, the excitatory peak was followed by an inhibitory dip occurring near stimulus offset, raising the possibility that excitation and inhibition jointly contributed to marking the onset and termination of the stimulus. This interpretation is conceptually relevant to recent work in moths showing that olfactory neurons can use excitatory and inhibitory phases to encode odor pulse timing and stimulus duration (Barta et al., 2024). Note that we did not systematically vary stimulus duration, and therefore do not directly demonstrate duration coding in the present study. Rather, our results suggest that onset/offset-related temporal structuring may also be present in central honey bee neurons responding to IPA.

Another important result of this study is that IPA-responsive activity was not restricted to the antennal lobe. We recorded two IPA-responsive protocerebral neurons. Both had branches spatially associated with mushroom body-related structures, the pedunculus, and surrounding protocerebral regions; one was also positioned near the anterior optic tubercle, antennal lobe, and adjacent protocerebral neuropils. These observations complement previous work on higher-order odor processing in the honey bee brain. Homberg showed that mushroom body-related extrinsic neurons can process antennal odor information, including isoamyl acetate, and can integrate it with mechanosensory, tactile, sucrose, and visual inputs (Homberg, 1984). Roussel et al. showed that alarm-related odors such as IPA and 2-heptanone evoke odor-specific activity patterns in the lateral horn (Roussel et al., 2014), and Carcaud et al. later showed, using GCaMP-expressing bees, that IPA evokes significant calcium responses in the lateral horn (Carcaud et al., 2023). Our protocerebral recordings show that some neurons beyond the primary olfactory center exhibited stimulus-related activity during IPA stimulation. Whether these protocerebral neurons respond selectively to odor input or also integrate other stimulus modalities, such as mechanosensory or visual signals, remains unresolved and will require dedicated control experiments.

These protocerebral results should nevertheless be interpreted cautiously. The number of successfully labeled and analyzed protocerebral neurons was limited, and their precise cell types, input sources, and downstream functions remain unresolved. Therefore, we cannot conclude that these neurons directly mediate IPA-induced defense, learning suppression, or foraging inhibition. A more conservative interpretation is that this study provides direct evidence that IPA can be represented by individual neurons outside the antennal lobe. Whether these protocerebral neurons participate in the integration of alarm information with internal state, prior experience, or other sensory cues will require a larger sample of identified neurons, more precise morphological classification, and behaviorally linked experiments.

Taken together, our results support a distributed and temporally dynamic model of IPA coding in the honey bee brain. Previous population-imaging studies have shown that IPA and other sting pheromone components evoke spatial activity patterns in the antennal lobe and lateral horn (Wang et al., 2008; Roussel et al., 2014; Carcaud et al., 2023). The present study suggests that these spatial patterns are accompanied by rich single-neuron temporal dynamics: different neurons show distinct response motifs, some neurons encode concentration through response magnitude and inhibition, and repeated stimulation can produce modest response attenuation and delayed timing. Rather than supporting the idea that IPA is read out by a single dedicated channel, our data favor a model in which IPA is represented across multiple neurons, temporal response patterns, and brain regions.

In summary, this study provides direct single-neuron evidence for central coding of the honey bee alarm pheromone component IPA. IPA did not evoke a uniform antennal lobe response; instead, it was represented by multiple temporal response motifs. IPA-responsive activity was also detected in protocerebral neurons, indicating that this alarm-related signal can be represented beyond the primary olfactory center. These findings extend previous behavioral and population-imaging studies to the single-neuron level and suggest that the honey bee brain represents IPA through distributed, dynamic, and hierarchical neural processes.

## CRediT authorship contribution statement

**Tao Lin**: Conceptualization, Methodology, Investigation, Formal analysis, Writing – original draft, Writing – review & editing, Funding acquisition. **Brian H. Smith**: Resources, Writing – review and editing. **Hong Lei**: Conceptualization, Methodology, Supervision, Resources, Writing – review & editing.

## Acknowledgments

This work was supported by Jiangxi Provincial Natural Science Foundation (Grant No. 20242BAB25352).

---

**Table S1.** Summary of representative kinetic response parameters for individual antennal lobe neurons. Response types were defined based on the temporal profile of the IPA-evoked response and were classified as fast transient, monophasic, biphasic excitation–inhibition (E– I), and delayed excitation–inhibition (E–I).

| Neuron ID | N trials | Response type | Peak amplitude (Hz) | Peak latency (s) | Response duration (s) | Negative AUC |
| --- | --- | --- | --- | --- | --- | --- |
| Neuron #1 | 5 | Fast transient | 83.60 | 0.28 | 0.050 | 0.42 |
| Neuron #2 | 4 | Monophasic | 20.50 | 0.38 | 0.25 | 5.08 |
| Neuron #3 | 6 | Biphasic E–I | 22.67 | 0.38 | 0.25 | 3.77 |
| Neuron #4 | 5 | Biphasic E–I | 38.40 | 0.48 | 0.30 | 1.68 |
| Neuron #5 | 5 | Monophasic | 20.80 | 0.32 | 0.70 | 1.12 |
| Neuron #6 | 5 | Monophasic | 24.40 | 0.58 | 0.85 | 3.42 |
| Neuron #7 | 5 | Biphasic E–I | 40.0 | 0.52 | 0.20 | 0.00 |
| Neuron #8 | 18 | Delayed E–I | 30.89 | 0.28 | 0.050 | 3.20 |

